# Road salt compromises functional morphology of larval gills in populations of an amphibian

**DOI:** 10.1101/2020.12.26.424459

**Authors:** Richard V. Szeligowski, Jules A. Scanley, Christine C. Broadbridge, Steven P. Brady

**Author notes:** Corresponding Author: Richard Szeligowski.

## Abstract

Throughout much of the world, winter deicing practices have led to secondary salinization of freshwater habitats, where numerous taxa are vulnerable to elevated salinity. Many amphibians are of particular concern because of their permeable skin and reliance on small ponds and pools, where salinity levels can be high. The early life-history stages of amphibians that develop in these habitats are especially sensitive to salt exposure. Larvae developing in salt-polluted environments must osmoregulate through ion exchange in gills. While salt-induced changes to the physiology of ion exchange in amphibian gills is generally understood, functionally relevant changes in gill morphology remain poorly described. Yet the structure of gills should be an important component affecting their ionoregulatory capacity, for instance in terms available surface area. Larval amphibian gills also play critical roles in gas exchange and foraging. Thus, changes in gill morphology due to salt pollution potentially affect not only osmoregulation, but also respiration and feeding. Here, we used a chronic exposure experiment to quantify the effect of salinity on larval gill morphology in populations of the wood frog (*Rana sylvatica*). We measured a suite of morphological traits on gill tufts, where ionoregulation and gas exchange occur, and on gill filters, which are used in feeding. Larvae raised in high salinity conditions had gill tufts with lower surface area to volume ratio, while epithelial cells on these tufts were less circular but occurred at higher densities. Gill filters showed increased spacing, which can potentially reduce their efficiency in filtering food particles. Together, these changes seem likely to diminish the ionoregulatory and respiratory capacity of gill tufts, and compromise feeding functionality of gill filters. Thus, a singular change in the aquatic environment from a widespread pollutant has the potential to generate a suite of consequences via changes in gill morphology. Critically, this suite of negative effects is likely most detrimental in salinized environments, where ionoregulatory demands are higher, which in turn should increase respiratory demands along with energy acquisition demands through foraging.

**Summary Statement:** Chronic road salt exposure alters the functional morphology of gills in larval amphibians, potentially compromising osmoregulation, feeding, and respiration.

## Introduction

### Salt pollution: a widespread problem for freshwater organisms

Freshwater habitats are becoming salinized by pollution from human activities including industrial processes, mining effluent, agricultural runoff, and most substantially, road salt used for de-icing (Lauer et al. 2016; Kaushal et al., 2005; Dugan et al., 2017; Maloney et al., 2017). Road salt applied to roadways, driveways, walkways, and parking lots runs off into adjacent habitats and/or stormwater infrastructure, and much of this salt is carried into surface waters. For instance, one estimate suggests that about 70% of lakes in the North American Lakes Region are polluted by road salt, and that if current salting practices continue, chloride pollution levels in many lakes will exceed the EPA’s water quality criterion 2050 (Dugan et al., 2017).

Elevated salinity in freshwater habitats spurs a suite of negative effects across biological levels, from genetic to ecosystem (Hintz et al., 2017; Gibbons et al., 2018; Merrick and Searle, 2019; Venâncio et al., 2019). For many freshwater taxa, salt pollution causes numerous sublethal effects, including changes in foraging and anti-predator behavior (Hall et al., 2017); changes in life history traits (Faulkner et al., 2019); and increased malformations (Brady, 2013; Hopkins et al., 2013; Alam et al., 2020). Salt can also be directly toxic (Sanzo and Hecnar, 2006; Karraker et al., 2008; Mahrosh et al., 2014). Together, these impacts can lead to loss of genetic diversity, population declines, and changes in community composition (Van Meter and Swan, 2014; Castillo et al., 2018), which collectively can modify ecosystem function (Schuler et al., 2017). Salt pollution also affects biogeochemical cycles, and these changes can scale up through food webs (Judd et al., 2005). For instance, salt pollution increases nitrogen and carbon export in streams (Reviewed in Hintz and Relyea, 2019) while plant and zooplankton richness have both been shown to decline in freshwater habitats polluted by salt (Richburg et al., 2001; Gutierrez et al., 2018).

While many freshwater taxa are affected by salt pollution, amphibians are of particular concern in part because of ongoing global population declines (Green et al., 2020). Although a subset of amphibians is tolerant of saline conditions (Hopkins and Brodie, 2015), many are considered vulnerable to increased salinization because they have narrow osmotic ranges (Chinathamby et al., 2006; Sanzo and Hecnar, 2006; Brady, 2013), possess highly permeable skin (Pough, 2007), and rely on freshwater habitats for reproduction, development, and dwelling (Vences and Köhler, 2008). While even modest increases in salt can be detrimental (Venâncio et al., 2019), concentrations can become especially high in shallow and/or temporary habitats (e.g., vernal pools), which many amphibians require to complete their lifecycle (Semlitsch and Skelly, 2007). Consequences of exposure to secondary salinization in amphibians include osmotic stress (Gomez-Mestre et al., 2004) and increased energy allocated to osmoregulation (Peña-Villalobos et al., 2016); increased risk of predation (Denoël et al., 2010); behavioral changes including reduced swim performance (Squires et al., 2008) and reduced foraging activity (Hall et al., 2017); decreased abundance (Albecker and McCoy, 2017); lower larval survival rates and ultimately, population size (Karraker et al., 2008).

### Osmoregulation in amphibians

Many amhibians have complex life cycles (Semlitsch and Skelly, 2007), and the negative effects of salt exposure tend to be most severe during early stages of development as embryos and larvae (Gomez-Mestre et al., 2004; Alexander et al., 2012; C.S. Wu et al., 2014; Park and Do, 2020). In anuran larvae, gills are the primary site of ionoregulation—a key mode of osmoregulation in which ions are exchanged between internal and external environments to maintain internal ion concentrations, especially Na^+^ and Cl^−^ (Alvarado and Moody, 1970; Dietz and Alvarado, 1974). For amphibians in typical (unpolluted) freshwater habitats, the tendency is for ions to be lost to the external environment. Osmoregulation for amphibians in these environments requires actively pumping ions in through gill epithelium into systemic plasma (Boonkoom and Alvarado, 1971). The primary mechanism responsible for ion exchange is the Na^+^/K^+^ ATPase ion pump (NKA). NKA is a transmembrane pump found in all animals that maintains body fluid homeostasis by exporting 3 Na^+^ ions while importing 2 K^+^ ions per ATP molecule against their respective concentration gradients (Skou, 1957; Pivovarov et al., 2019; Fig. 1). NKA-mediated osmoregulation operates via passive influx of Na^+^ from the external environment into an apical epithelial cell. From there, NKA actively transports Na^+^ ions into the Na^+^-rich plasma while exchanging K^+^ from the plasma into the cytosol, where K^+^ concentrations are higher. K^+^ ions exit the cytosol back into the plasma via passive membrane channels, which creates an electric potential between the external environment and the plasma, leading to passive influx of Cl^−^ ions and creating a hyperosmotic internal environment (Campbell et al., 2012).

**Figure 1.**
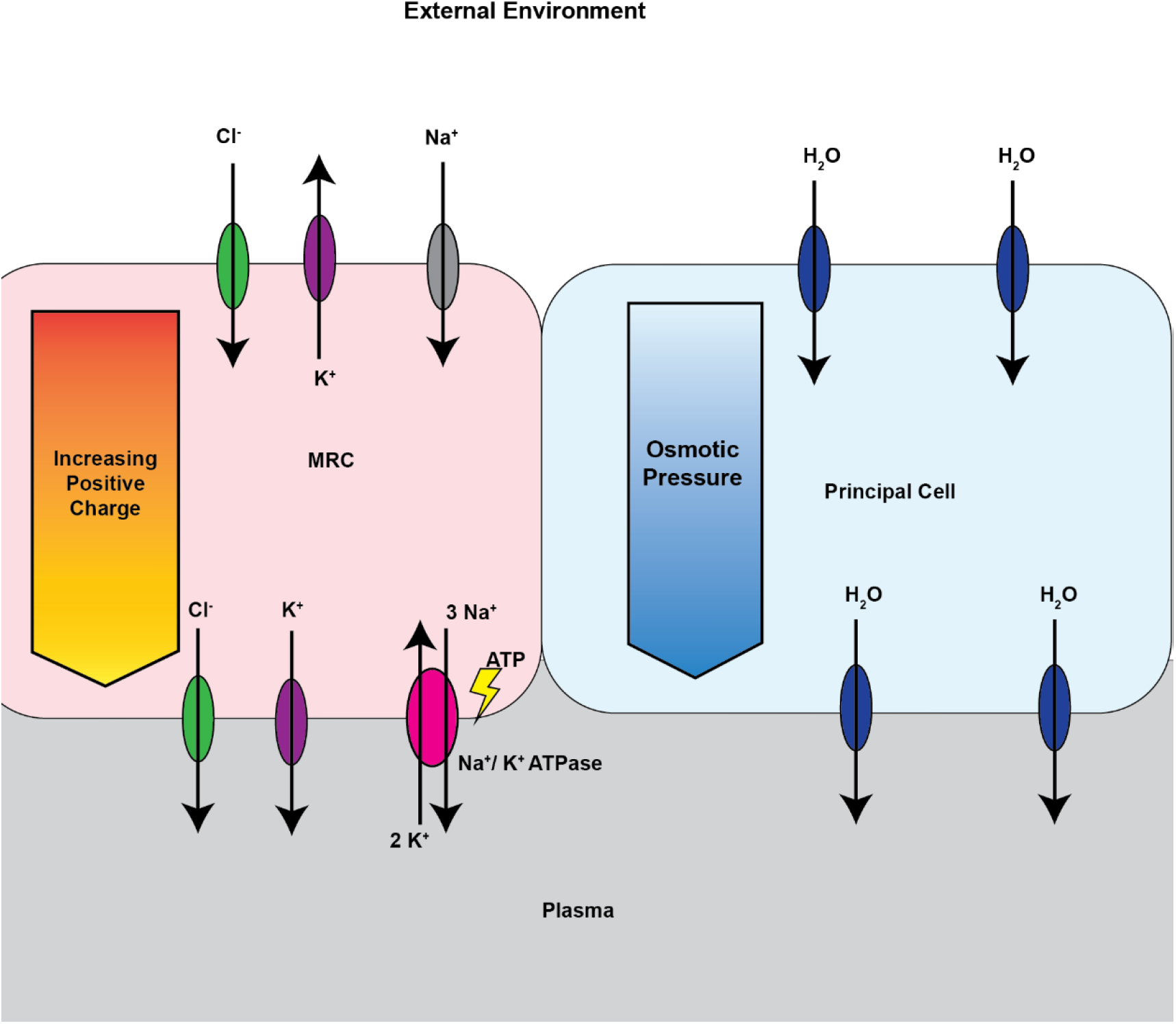
Ionic and Osmotic Control of Osmoregulation Across Gill Epithelium. Mitochondria rich cells (MRC) create a gradient of increasing positive charge through the action of Na^+^/K^+^ ATPase (NKA) pumps in the basal cell membrane. 3 Na^+^ ions are actively pumped out of the cell and 2 K^+^ ions are pumped in for every molecule of ATP. This creates a chemical gradient for Na^+^ to be drawn into the cytosol through passive Na^+^ channels. K^+^ is present in higher concentrations inside the cell and so passively exits through K^+^ channels into the plasma or the external environment. This high concentration of positive ions in the plasma generates a voltage gradient that allows Cl^−^ from the environment to pass through the cytosol into the plasma. The osmolality of the plasma is thus increased and aquaporins present in principle cells allow the passive diffusion of water.

Anuran gills are thought to function much like those of teleosts (Ultsch et al., 1999) in that NKA is highly expressed in mitochondrion-rich cells (MRCs) in the gill epithelium, where MRCs function as the primary mechanism of ionoregulation (Hwang and Lee, 2007). When freshwater amphibian species experience osmoregulatory challenges through salinization, they must prevent water loss by increasing ion absorption, thereby maintaining a hyperosmotic internal environment (Gomez-Mestre et al., 2004). Bernabò et al. (2013) found that larvae of the common toad (*Bufo bufo*) and Balearic green toad (*Bufo balearicus*) increased NKA abundance in response to increasing salinity, presumably to maintain a hyperosmotic internal environment and prevent water loss. Euryhaline (i.e., salinity tolerant) amphibians such as the Indian rice frog (*Fejervarya limnocharis*) seem to utilize the same strategy for osmoregulation, although the specific mechanisms that allow increased salinity tolerance are not fully understood. Wu *et al.* (2014) found that crab-eating frog (*Rana cancrivora*) tadpoles acclimated to salinity levels corresponding to 15,700 μS/cm had a six-fold increase in NKA abundance when compared to those raised in 4,000 μS/cm water. Although crab-eating frog tadpoles can survive in full strength seawater after acclimation (Gordon and Tucker, 1965), Hsu *et al.* (2012) found that metamorphosis occurred in only 40% of tadpoles in salinity levels corresponding to 60% seawater (30,000 μS/cm) after salinity acclimation.

### Toward improved understanding of gill morphology in osmoregulation

In contrast to our knowledge of *physiological* mechanisms underpinning osmoregulation in larval amphibians, we have limited understanding of the functional role of gill morphology, and how it might be affected by salt exposure. Yet changes in gill morphology are important adaptations in fish that transition between freshwater and seawater (Uchida et al., 1996; Hiroi et al., 1999), and thus might be important in freshwater amphibians exposed to salt. Limited, primarily qualitative insights from an acute exposure experiment show that high salinity can modify gill morphology in larvae of the green toad (*Bufo balearicus*) and common toad (*Bufo bufo*) (Bernabò et al., 2013). Specifically, gill tufts became “dehydrated” and “invaginated” in larvae exposed to 12,700 μS/cm. For larvae exposed to 15,000 μS/cm, hyperplasia was observed in pavement cells, which play a limited role in ion exchange (Wilson, 2011). While these qualitative insights are important, they were based on short-term exposure to salt concentrations far above those found in most secondarily-salinized freshwater habitats. Thus, a quantitative assessment of morphology, based on chronic exposure to ecologically relevant salinity levels for pollution is critically needed.

Changes in larval amphibian gill filter morphology could also affect gas exchange and therefore respiration. And because gills are also important structures for larval feeding, salt-induced morphological changes may hamper foraging efficiency. For instance, if gill morphology becomes less well suited to trapping food particles, larvae should experience relative increases in foraging demand. Conceivably, these various effects could be multiplicative: increased demands for ionoregulation in salt-polluted environments could change gill morphology in ways that reduce efficiency for ionoregulation and gas exchange, driving energetic demands higher. At the same time, reduced foraging efficiency could further increase energetic demands. Thus, any changes in gill morphology have the potential to generate a cascade of effects on amphibian physiology and fitness.

Here, we begin exploring these potential effects by quantifying changes in gill morphology in populations of the wood frog (*Rana sylvatica*). We raised wild-collected embryos in low and high salinity conditions, from early embryonic stages until nearing metamorphosis. We used scanning electron microscopy to image and quantify changes in gill morphology in response to salinity. We hypothesized that increased salinity would alter gill morphology, particularly at the gill tufts and filters. These sites, respectively, are where most osmoregulation occurs via NKA activity (Brunelli et al., 2004), and where food particles are filtered for feeding (Seale and Wassersug, 1979). We predicted that salt would modify gill morphology in ways that compromise ionoregulatory capacity, gas exchange, and feeding.

## Methods

### Natural History and larval gill function

Wood frogs are found throughout the northern ranges of North America, from the southern Appalachian Mountains through Canadian and Alaskan forests in habitats including tundra, deciduous and coniferous forests, marshes, and grasslands (Green et al., 2014). Wood frog breeding is seasonal. In our study region in the northeastern U.S., adults migrate from upland terrestrial habitat to nearby small, shallow, and often temporary ponds, typically during the first warm rain of early spring following hibernation. Breeding is generally explosive, occurring over a 1-2 week period in each pond. Each female lays one clutch as a single egg masses containing about 1,000 eggs, which are fertilized externally during amplexus. Eggs develop for about 2-3 weeks before hatching, after which time larvae (i.e., tadpoles) develop for about two months before metamorphosing into terrestrial juveniles.

Among amphibians, the wood frog stands out as having a particularly puzzling response to salt exposure. Not surprisingly, this species shows a variety of negative responses to salt pollution and roadside dwelling including many of the known negative effects associated with salt exposure, such as direct mortality and increased malformation in embryos and larvae (Karraker et al., 2008; Brady, 2013) as well as reduced growth rates and increased developmental variance (Brady, 2013; Dananay et al., 2015). However, reciprocal transplants and common garden experiments reveal that aquatic-stage wood frogs from salt-polluted roadside ponds are *more* sensitive to road salt and roadside conditions than nearby populations from unpolluted ponds (Brady, 2013; 2017). These findings suggest that roadside populations might have recently evolved lower tolerance to salt than individuals from unpolluted populations, consistent with ‘local maladaptation’ processes (Brady et al., 2019). Alternatively, this pattern might reflect a fitness tradeoff between aquatic- and terrestrial-stage frogs, for instance if lower salt tolerance is correlated with adult lifetime reproductive success. In either case, we speculate that the survival disadvantage in roadside populations might arise through effects on gill morphology.

During larval development, the primary site of respiration changes. Prior to feeding stage - Gosner (Gosner, 1960) stage 25 - respiration occurs primarily via cutaneous diffusion and the limited activity of external gills before transitioning to gas exchange in a bilateral pair of internal gills (Feder and Burggren, 1992; Brunelli, 2004). Each internal gill contains three prominent branchial arches that originate in the rostral end of the gill, extending caudally. A reduced fourth branchial arch lateral to the first three arches is present in some species but has decreased tuft density and filter plate size (McIndoe and Smith, 1984; Brunelli et al., 2004). Gill tufts are the primary site of gas exchange (McIndoe and Smith, 1984; Brunelli et al., 2004), protruding ventrally from the branchial arch. Each tuft has numerous smaller structures that split from the main body (hereafter ‘tuft branches’). A cartilaginous filter plate protrudes dorsally from the branchial arch (McIndoe and Smith, 1984), and rows of gill filters extend laterally from the filter plate, with main folds running ventrodorsally in a zigzag pattern (McIndoe and Smith, 1984). Secondary folds branch from the main folds at corners, often continuing the zigzag pattern of main folds, tertiary folds branch away from secondary folds at their corners, often forming a rectangular filter niche with the main and secondary folds (McIndoe and Smith, 1984). We termed points where folds changed directions, branched into other folds, or terminated ‘nodes’.

Gill filters function as a screen for food particulates as water travels over them towards the tufts (Feder et al., 1984; McIndoe and Smith, 1984; Brunelli et al., 2004). Both gill tufts and filters play roles in ionoregulation. However, tufts seem to play a greater part than filters in this capacity. For instance, Bernabò et al. (2013) showed NKA is more strongly expressed in gill tufts than filters in both common toad and Balearic green toad gills, although NKA expression increased in tufts and filters for both species when exposed to salt levels between 5%-25% sea water.

### Embryo collection and rearing

We collected egg masses from six natural ponds in Norwich, VT during the breeding season in April 2019. To represent a range of populations, we sampled egg masses from three salt-polluted ‘roadside’ ponds located within 10 m of a paved road and three unpolluted ‘woodland’ ponds, which were located at least 500 m from any road. From each pond, a cluster of approximately 20 embryos were removed from each of five egg masses. Selection of egg masses and embryo groups was haphazard. Embryo clusters were placed in plastic 700 ml (16.5 × 7.5 × 12.5 cm) containers filled with natal pond water and were immediately transported to the lab in New Haven, CT where they were stored at 4°C for 24 hours until the start of the chronic salt-exposure experiment.

Four salinity treatments were prepared using aged, conditioned tap water and NaCl road salt obtained from the Connecticut Department of Transportation. Embryos were exposed to one of four treatments: 275 μS/cm (no salt added), 1,000 μS/cm, 4,000 μS/cm, and 7,000 μS/cm. These values were chosen, respectively, to represent control conditions, average roadside conditions, maximum roadside conditions, and values exceeding roadside conditions to consider a broad range of exposure scenarios. Embryo clusters were assigned haphazardly to one of four salinity treatments in a 5.6 L container (30.5 × 10 × 20 cm) containing 5 L of assigned treatment solution such that each pond had representative embryos raised in each treatment solution. Containers were arranged randomly across four levels of a shelving unit and were rotated weekly within and between shelves to distribute potential block effects. When tadpoles reached feeding stage, they were fed 5-day food rations containing a 3:1 ratio of rabbit chow to fish food concurrently with water changes. Tadpoles were raised until they reached approximately Gosner stage 35. After recording tadpole developmental stage scores, individuals were euthanized via immersion in buffered Tricaine (2g/L MS222 ethyl 3-aminobenzoate methanesulfonate, Sigma-Aldrich; pH 7.0) and gills were immediately excised and prepped for scanning electron microscopy (SEM). All methods complied with institutional animal care guidelines

### Scanning electron microscopy sample preparation

We selected tadpoles haphazardly for SEM, imaging 19 individual tadpoles representing different salt treatments and populations (see Supp Table 1). This sample size was chosen to balance SEM constraints (e.g., costs, sample prep time, personnel time) with study goals. Selected individuals ranged in Gosner developmental stage from 34-37. For each tadpole, the left gill was excised, measured for length using the reticle of a dissecting scope, and immediately transferred to 3% glutaraldehyde in phosphate buffer at 4°C for 1.5 hours, then transferred into 1% osmium tetroxide in phosphate buffer for 2 hours at 4°C. Gills were then dehydrated in a series of graded ethanol steps (30%, 50%, 70%, 80%, 90% and 100%), modified from Brunelli et. al (2004). Gills were dried in a Tousimis Samdri-780 critical point drier and fixed to SEM stubs. Gills were imaged with a Zeiss Sigma VP field emission scanning electron microscope.

### Gill Morphology

Images were analyzed using ImageJ software (v. 1.51, Schneider et al. 2012). To describe potential effects of salt on gill morphology, we targeted measuring 12 different traits per gill (Table 1). We measured gill tuft traits (N=7) corresponding to available surface area (e.g. tuft branch length and width), gill filter traits (N=5) corresponding with surface area available for ion exchange (e.g. internode length), and traits related to filtering efficacy (e.g. internode length and angle). Tuft branch cell attributes were also measured to assess how cell morphology may contribute to ultrastructure (observable through electron microscopy) changes. Each gill was assessed for multiple traits; however, SEM imaging could not guarantee comprehensive coverage of all traits on each gill (see Supplemental Table 1).

**Table 1.** Description of gill traits measured in response to salinity.

| Gill Trait | Description |
| --- | --- |
| **Gill Tufts** |  |
| *Branch Length ($T_L$)* | Length of tuft branch from base of branch to tip. Each datapoint for this trait in our study is the average of three such measurements per branch to account for imprecise branch origins. |
| *Branch Width ($T_W$)* | Width of tuft branch. Each datapoint for this trait in our study is the average of three such measurements per branch to account for tapering of branches from broader bases to narrower termini. |
| *Cell Area* | Area of gill tuft cells as defined by cell borders at apical epithelium. |
| *Cell Perimeter* | Perimeter of gill tuft cells as defined by cell borders at apical epithelium. |
| *Cell Aspect Ratio* | Ratio of the major axis, or longest diameter through the cell, to the minor axis, or shortest diameter through the cell. |
| *Cell Circularity* | A measure of how closely the perimeter of the cell describes a circle. Given by the formula<br>$4\pi \frac{\text{Area of ellipse}}{\text{perimeter of ellipse}^2}$<br>A circularity value of 1 denotes a perfect circle. |
| *Cell Density* | The number of cells present within tuft cell branches. |
| **Gill Filters** |  |
| *Main Fold Spacing* | The distance between rows of main folds. Multiple measures were taken between folds to account for differences arising from the zigzagging morphology of gill filter folds. |
| *Main Fold Internode Distance* | The distance between points in main folds where either the main fold changed direction, a secondary fold branched from the main fold, or the main fold terminated. |
| *Main Fold to Main Fold Angle* | The angle described by two adjacent sections of main fold, joined by a node, that do not lie on the same line |
| *Secondary Fold Internode Distance* | The distance between points on secondary folds where either the secondary node joined the main fold, changed direction, a tertiary fold branched from the secondary fold, or the secondary fold terminated. |
| *Main Fold to Secondary Fold Angle* | The angle described by two adjacent sections of main and secondary folds of the gill filter. Angles were always measured from the ventral side of the main fold onto the secondary fold. |

For gill tufts, we measured epithelial cell attributes including cell area (C_A_) and cell perimeter (C_P_) by tracing the perimeter of cells where boundaries were clearly definable, targeting 25 cells per gill (illustrated in Fig. 2a). Cell aspect ratio and circularity were calculated from these values to characterize cell shape. Tuft cell density (T_D_) was calculated by tracing clusters of cells within tuft branches and counting the number of cells in each defined area, then computing the value of cell count over area. Tuft branch measurements were obtained by averaging three linear measurements in tuft branch length (T_L_) and tuft branch width (T_W_), respectively (illustrated in Fig. 3a). We averaged three measurements per trait per branch to account for nonuniform width along branches and imprecise branch origins.

**Figure 2.**
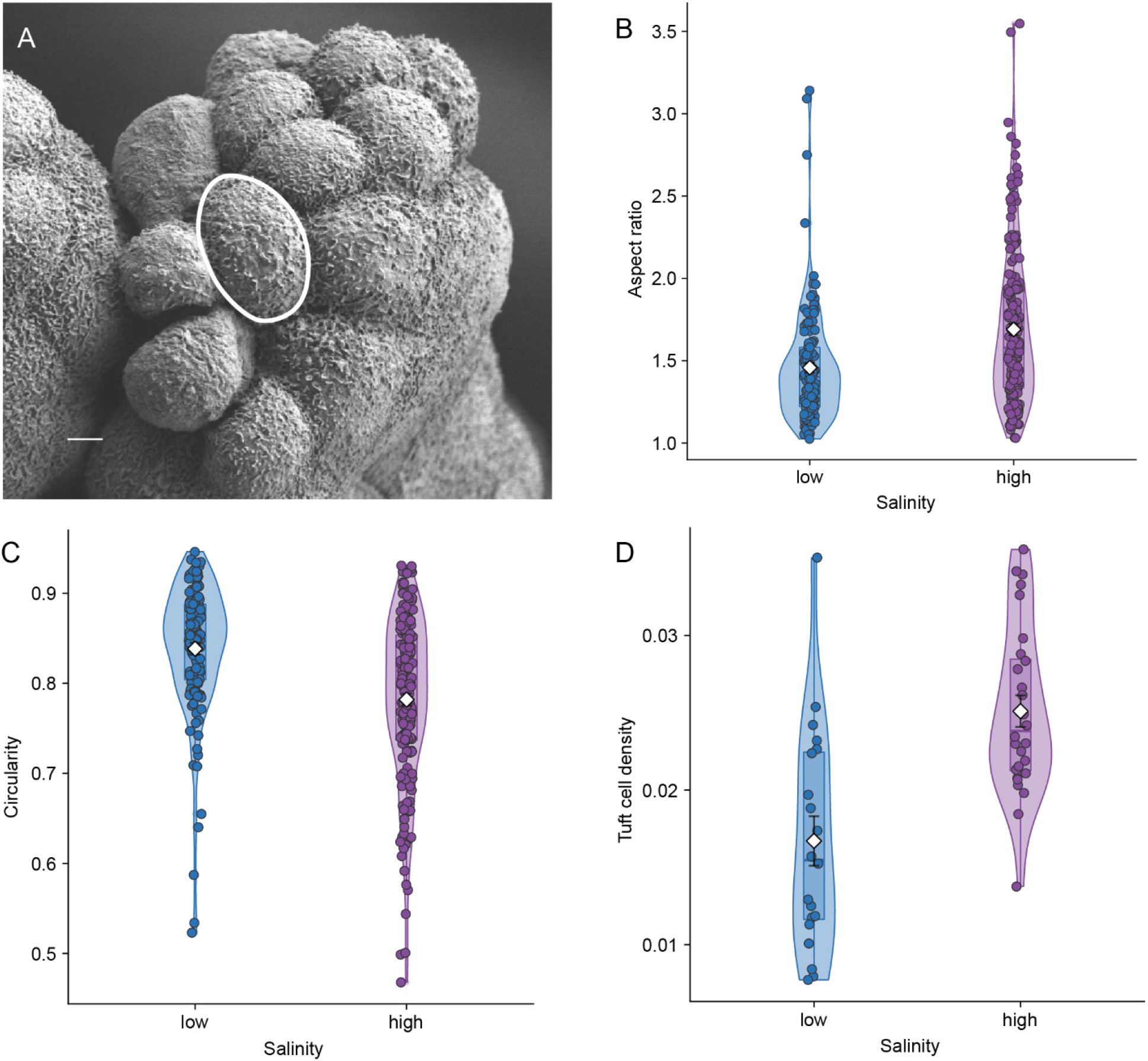
Variation in Tuft Branch Cell Shape and Density. Gill tuft branch at 7000x magnification (A) with a tuft cell circled in white. Scale bar, 2 μm. Cell shape was measured by tracing cells with well-defined borders and measuring area, perimeter, aspect ratio (B), and circularity (C). Tuft cell density (D) was obtained by tracing cell branches and counting cells within measured area. Points represent individual measurements and were jittered to reduce overlap; violin plots show density distributions, overlayed with a boxplot. Means with SE bars are shown with white diamond. Aspect ratio increased with salinity and circularity decreased. Altered cell shape in high salt treatments coincided with higher cell density.

**Figure 3.**
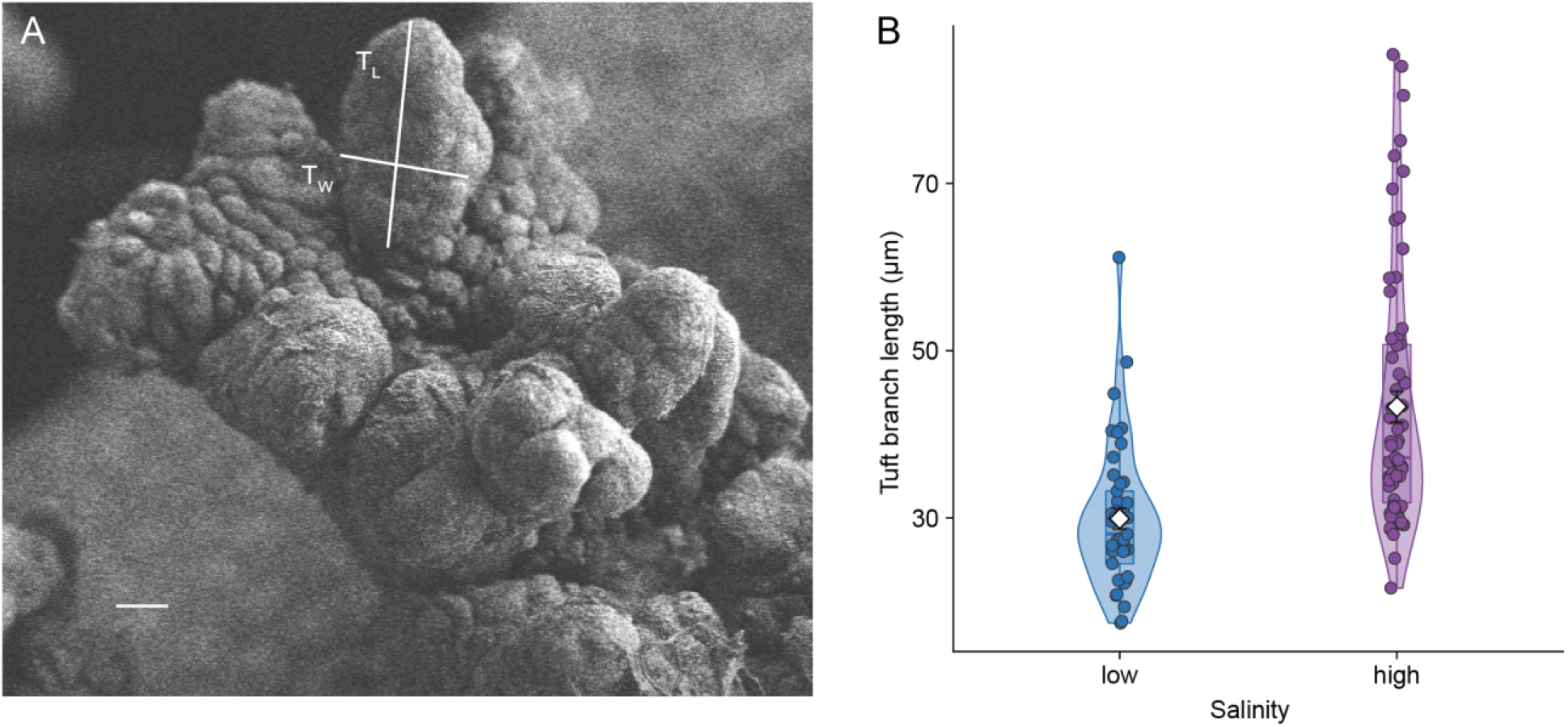
Gill Tuft Branch Variation in response to salt. Gill tuft (A) with representative tuft branch length (T_L_) and tuft branch width (T_W_) depicted in white. Scale bar, 10 μm. Each datapoint for tuft dimensions is the average of three measurements taken per trait per tuft branch. Tuft branches are nonuniform in width, measurements were taken at towards the base, midsection, and apex accordingly. Tuft branch length (B) was averaged due to the boundary of the tuft branch at its base being imprecise. Points represent individual measurements and were jittered to reduce overlap; violin plots show density distributions, overlayed with a boxplot. Means with SE bars are shown with white diamond. T_L_ increased in response to salt, was less densely distributed around the mean and had a wider range of values. T_W_ was not found to vary in response to salt.

For gill filters, we measured internode distance for both main folds (D_M_) and secondary folds (D_S_) (Fig. 4a). We also measured angles between main folds (φ_MM_) and secondary folds (φ_MS_) (Fig. 4a). Measurements for φ_MS_ angles were measured starting from the ventral end of the main fold (proximal to branchial arch) and tracing the angle of the secondary fold, φ_MM_ angles were measured ventrodorsally. The distance between main folds (D_MM_) on filter plates (Fig. 4a) was measured by taking the average of three measurements of the distance from a point on the main fold apparatus (corresponding to nodes and points midway between nodes) to the nearest point on an adjacent main fold apparatus.

**Figure 4.**
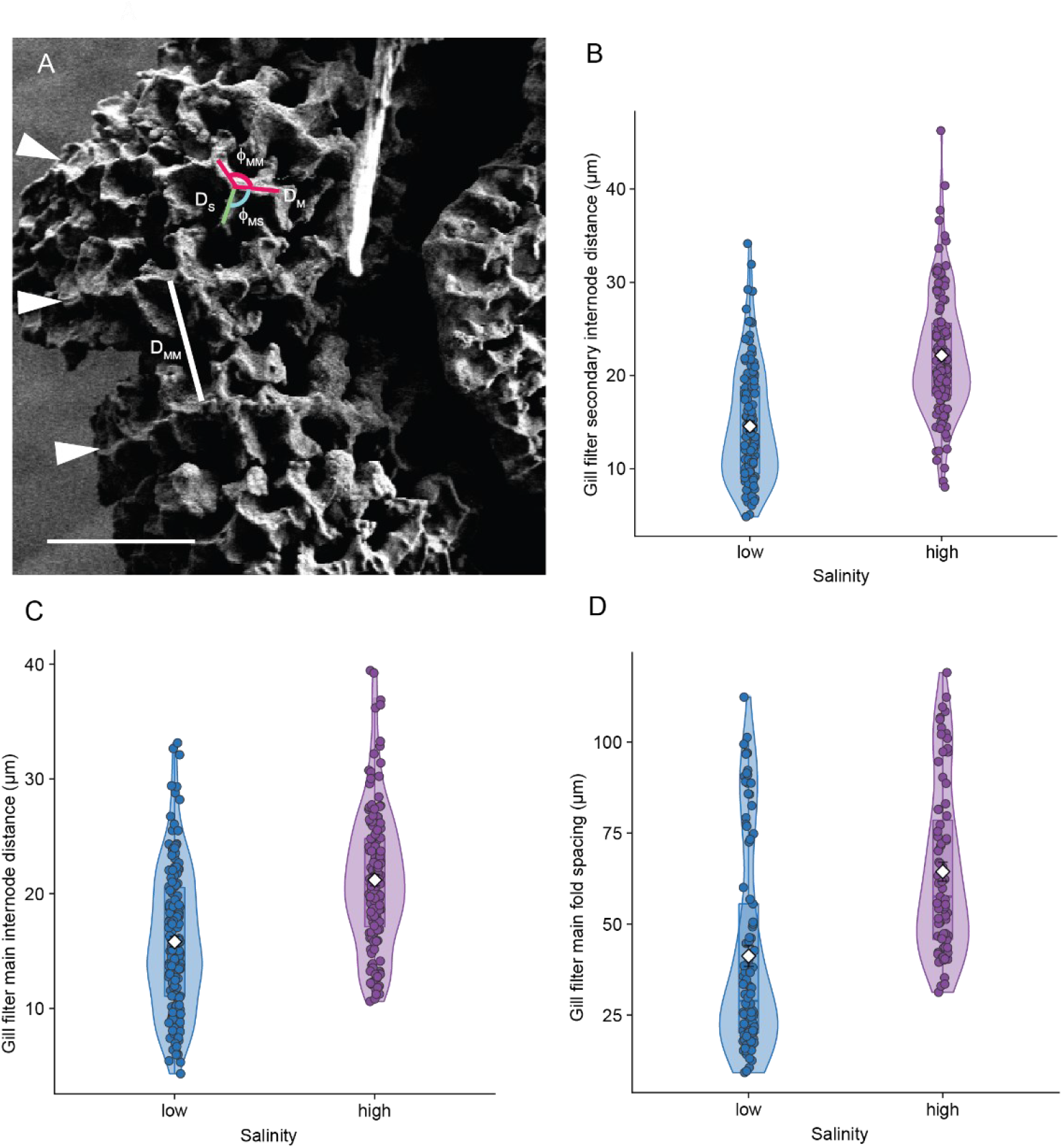
Gill Filter Fold Variation in Response to Salt. Gill plate with filters of epithelial tissue exposed (A). Scale bar, 100 μm. Main folds (arrows) run ventrodorsally along the gill plate (right to left in image above). Representative main internode distances (D_M_) are depicted in red, and main fold angle (φ_MM_) is represented by the red arc between lines. A representative secondary internode distance (D_S_) is depicted in green, and the main fold-secondary fold angle (φ_MS_) is depicted by the blue arc. A representative measurement of the distance between main fold rows (D_MM_) is shown in white. D_S_ (B), D_M_ (C), and D_MM_ (D) distances. Points represent individual measurements and were jittered to reduce overlap; violin plots show density distributions, overlayed with a boxplot. Means with SE bars are shown with white diamond. D_S_ increased in response to salinity.

**Figure 5.**
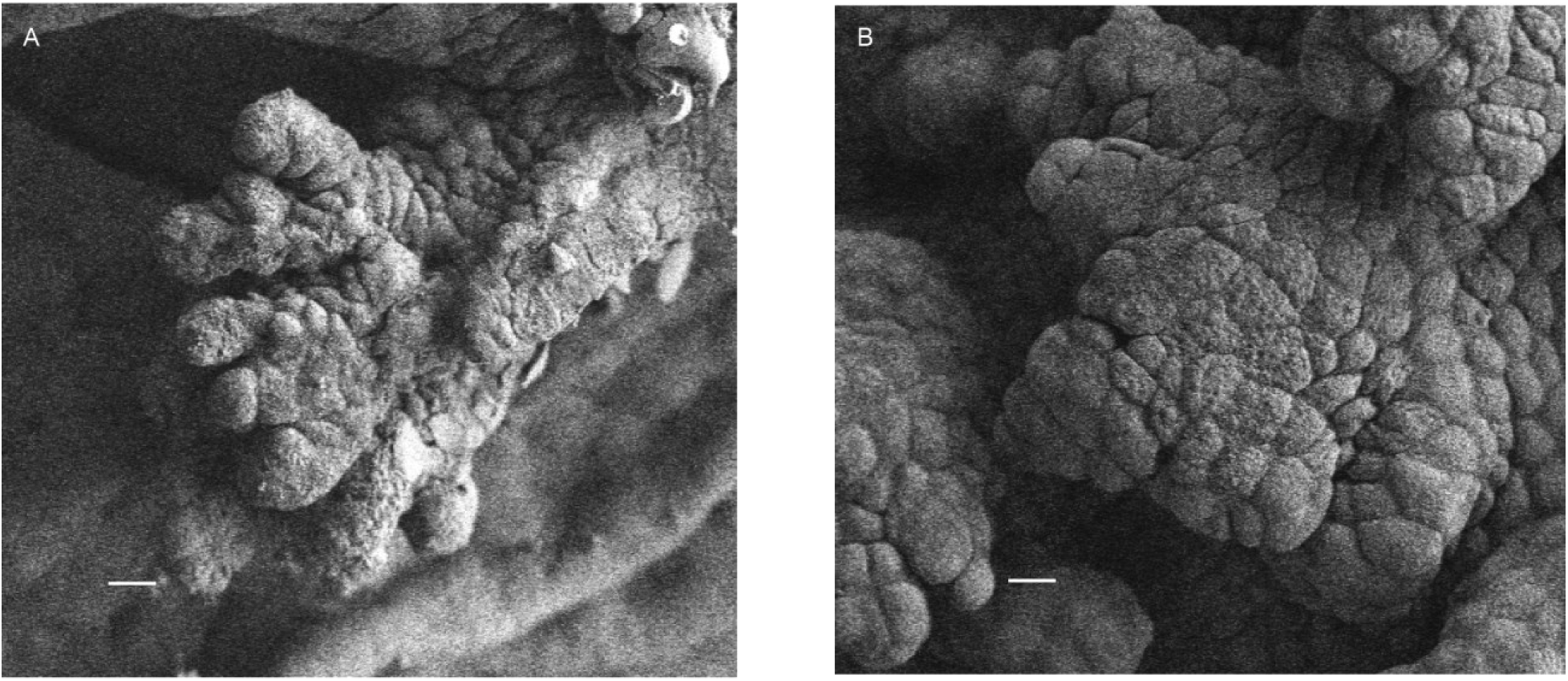
Comparison of Gill Tufts from Control and Salt Treatments. (A) Tufts from animals in control treatments were contracted relative to the fuller appearance of (B) tufts from salt treatments. Scale bars,10 μm.

### Analysis

To maximize power given our sample size, we composited data into two groups instead of the original four treatment groups – ‘high salinity’ comprised the three different salt concentrations while ‘low salinity’ comprised the control group. We used the ‘lme4’ package (Haubo Bojesen et al., 2017) in RStudio (v. 3.6.2, RStudio Team, 2020) to compose linear mixed effects models analyzing each gill trait in response to salinity. We included gill length and developmental stage as fixed effect covariates because of their potential to influence morphology. In each model, pond and gill identity were included as random effects to account for correlation within ponds and among repeated measurements on each gill. For each trait, we used AIC selection to choose the preferred random effects structure with a delta AIC threshold value of 2. Initial models were fit for AIC selection using restricted maximum likelihood, and selected models for inference were refit with maximum likelihood. *P*-values for fixed effects were based on degrees of freedom calculated with Satterthwaite’s approach implemented in the R package ‘lmerTest’ (Haubo Bojesen et al., 2017). We estimated surface area and volume for gill tuft branches by treating them as idealized cylinders and used model estimates for T_L_. Because T_W_ did not vary significantly, the average of T_W_ estimates was used for surface area and volume estimates. We estimated the filter niche area (i.e., size of spaces within filters) by treating them as rectangles with sides D_M_ and D_S_ and used model estimates for D_S_. As D_M_ did not vary significantly, we used the average of D_M_ model estimates for low and high salinity treatments.

## Results

Tuft branches were longer in high versus low salinity treatments (*F*_*1, 7.4*_ = 8.3, *P* = 0.022; fig. 3b). Specifically, model estimates for tuft branch length were 31.6 μm for gills in the low salinity treatments compared to 47.2 μm in the high salinity treatments. Thus, gill tuft branch length increased by an estimated 49.5% in tadpoles raised in high salinity conditions. Gill tuft branch width (*F*_*1, 9.4*_ = 0.5, *P* = 0.484) did not vary between salinity treatments. Calculations of branch surface area and volume using these model estimates gave a 41.86% increase in surface area from low salinity to high salinity treatments, but a 49.46% increase in tuft branch volume, reducing the surface area to volume ratio by 5.4% in high salinity treatments.

Traits associated with tuft cells also varied in response to salinity: cell aspect ratio (*F*_*1, 294*_ = 15.6, *P* = 0.001; fig. 2b), cell circularity (*F*_*1, 5.0*_ = 13.1, *P* = 0.0152; fig. 2c) and cell density (*F*_*1, 9.9*_ = 7.0, *P* = 0.025; fig. 2d) each varied in response to salinity. Model estimates for aspect ratio was 1.45 in low salinity treatments versus 1.63 in high salinity treatments, representing an 11.9% increase. Circularity in low salinity was estimated to be 0.611 compared to 0.562 in high salinity, revealing a 7.96% decrease in circularity for high salinity treatments. Cell density was 47.4% higher in high salinity tadpoles, with 0.025 cells per μm^2^ compared to 0.017 cells per μm^2^ in low salinity treatments. Tuft cell perimeter (*F*_*1, 8.0*_ = 1.5, *P* = 0.260) and tuft cell area (*F*_*1, 8.0*_ = 1.0, *P* = 0.354). No significant variation in any tuft trait occurred in response to Gosner developmental stage or gill length (all *P* > 0.166).

The distance between gill filter secondary nodes (D_S_) varied between low and high salinity treatments (*F*_*1, 4.8*_ = 10.3, *P* = 0.026; fig. 4b). In low salinity treatments, D_S_ was estimated to be 13.52 μm compared to 22.96 μm for high salinity treatments, representing a 68.52% increase in D_S_ for tadpoles raised in high salinity. There was marginal support for an effect of salinity on main fold internode distance (D_M_) (*F*_*1, 4.8*_ = 6.0, *P* = 0.060; fig. 4c) and distance between main folds (D_MM_) (*F*_*1, 4.0*_ = 5.4, *P* = 0.081; fig. 4d). Specifically, D_M_ model estimates were 14.43 μm in low salinity treatments and 21.34 μm in high salinity, representing a 32.36% increase in D_M_ in high salinity. D_MM_ model estimates were 45.76 μm in low salinity treatments and 79.26 μm in high salinity treatments, representing a 57.74% increase in high salinity. Main fold-main fold angles (φ_MM_) (*F*_*1, 4.3*_ = 4.0, *P* = 0.111), and main fold-secondary fold angles (φ_MS_) (*F*_*1, 3.9*_ = 0.8, *P* = 0.431) did not vary significantly between animals raised in low salinity and high salinity conditions. Calculating changes in filter niche area by using model estimates gave a 69.23% increase for filter niche area in high salinity. While neither Gosner developmental stage nor gill length covariates were correlated with filter trait variation in most of these models (D_M_, D_S_, φ_MM_, φ_MS_; *P* > 0.411), D_MM_ was correlated with gill length (*F*_*1, 4.0*_ = 5.4, *P* > 0.060), increasing 25.9 μm with each 1 mm of gill length.

## Discussion

High salinity had numerous effects on tadpole gill morphology. Tuft branch length increased by nearly half while its width was unchanged. Cells on these tufts were more tightly packed together and were less circular, with relatively longer major than minor axes, but cell perimeter and area were unchanged. For gill filters, the distance between nodes was longer, which should result in larger filter size. Together, these results demonstrate that chronic exposure to road salt modifies gill morphology in ways that reduce available surface area to volume ratio and increase the spacing of gill filters. These changes appear likely to compromise the functional capacity of gills for osmoregulation, gas exchange, and foraging.

Gill tufts are the primary site of ionoregulation and gas exchange. Increased tuft length (T_L_) in the absence of increased width (T_W_), strongly suggests that changes in morphology due to salt reduce the available surface area to volume ratio, thus reducing the efficiency and capacity of ion and gas exchange. Indeed, we estimated a 5.4% reduction in surface area to volume ratio for larvae from high salinity treatments. In addition to presumably lowering the efficiency of ion exchange, this change should also reduce the efficiency of gas exchange, which in turn likely exacerbates the already-increased metabolic demands of elevated salinity (Peña-Villalobos et al. 2016). The increase in cell density on tufts was attributable to changes in cell shape, allowing tighter packing of cells on the gill tufts. An increase in cell density could be a response to increase the rate of ion exchange, as no difference in cell area was found in high salinity treatments. While this could offset the reduction in the surface area to volume ratio found in high salinity treatments, a higher cell density is likely to incur a higher energetic cost. However, more information about cell-specific metabolic rates is needed to determine the extent of this tradeoff.

Gill filters screen food and particulates out of the water prior to gas exchange at the tufts, and thus play an important role in foraging and feeding (Feder et al. 1984; McIndoe and Smith 1984; Brunelli et al. 2004). Here, filter size appeared to increase in response to salinity, as evidenced by increased length of secondary node distance and (cautiously) the marginal evidence for increased distance between main folds and main fold internode distance. Increased filter size presumably reduces the efficacy of filtering food particles, as more food particles are likely to be flushed out through the operculum instead of being retained for consumption (Seale and Wassersug, 1979). In turn, foraging efficiency should decrease due to high salinity, and this particular change in morphology could bear an energetic cost as larvae might be required to increase foraging efforts. However, this change in filter structure likely increases the available area for ion exchange along the matrix (so-called ‘ramifications’) of the gill filters. Thus, there might be a tradeoff between foraging efficacy and osmoregulatory capacity.

A reduction in the ability of gill filters to both capture food and remove particulates from incoming water before it reaches the tufts may have compounding effects, leading to the widely reported lethal and sublethal consequences of salinity exposure on amphibians. Allowing larger particles to contact and potenitally entangle within the tufts (dowstream of the filters) could reduce the available area for gas exchange. This effect would be magnified by increases in NKA expression in response to salt (Bernabò et al., 2013; Wu et al., 2014) and the associated increase in the metabolic demands of osmoregulation (Peña-Villalobos et al., 2016). When these physiological effects are considered along with observed reductions in foraging activity (Hall et al., 2017), a decrease in the filter’s ability to capture food could have substantial impacts on survival. Diminshed feeding capacity coupled with larger, less efficient gill tufts likely constrains larval metabolic resources that are necessary to cope with the osmoregulatory burden of salt-polluted water.

It is interesting to consider that even though gill tuft surface area to volume ratio declined in response to high salinity, the length of gill tuft branches and filter secondary folds increased. This pattern of increased size contrasts with other morphological trait responses for amphibians exposed to high salinity. For instance, body length and mass have been shown to decrease in response to salt in numerous studies (e.g., Gomez-Mestre and Tejedo, 2003; Wu and Kam, 2009). Thus, the effect of high salinity on amphibian morphology appears to be tissue-specific. Osmoregulatory mechanisms in tadpole gill tissues might play a role in these differences via changes in voltage gradients across membranes. Non-excitable cell depolarization is associated with tissue growth and development (Shi and Borgens, 1995; Reid and Zhao, 2014), and the ionic gradient that tadpoles maintain across gill epithelium for osmoregulatory function creates a voltage gradient that Alvarado and Moody (1970) measured at −5 to −10 mV in pond water, which depolarized as salinity increased to 20mM/L (1,950 μS/cm). While further work is needed to characterize the mechanism behind increased gill tuft and filter size in response to salinity, we are intrigued by the possibility of salinity-induced depolarization contributing to the increases in gill structure size.

## Acknowledgements

We acknowledge the use of facilities and support (personnel and supplies) from the CT State Colleges and Universities Center for Nanotechnology, Werth Industry Academic Fellowship, and New Haven Innovation Collaborative Undergraduate Research Fellowship Programs. We thank E. Brunelli for providing expertise in amphibian gill anatomy. We are grateful for field and laboratory work by M. Forgione, L. Frymus, J. Priester, F. Senturk, and A. Tucker.

## Data Availability

All data will be made available in a public repository upon acceptance.

